# The aging musician: Evidence of a downward trend in song tempo as a function of artist age

**DOI:** 10.1101/2024.06.29.601154

**Authors:** Geoff Luck, Alessandro Ansani

## Abstract

Correspondences between the timing of motor behavior and that of musical performance are well-established. Motor behavior, however, is known to degrade across the adult lifespan due to neurobiological decay. In particular, performance on speed-dependent motor tasks deteriorates, spontaneous motor tempo (SMT) slows, and upper motor rate limit falls. Here, we examine whether this slowdown in motor behavior impacts tempo of musical performance as a function of age. We analyzed 14,556 songs released between 1956 and 2020 by artists with careers spanning at least 20 years. Generalized Additive Mixed Models (GAMMs) and Linear Mixed Models (LMMs) were employed to assess the effects of age, operationalized by subtracting birth year from release year of each track, on musical tempo. Results revealed a slight tempo increase from early adulthood to age 30, followed by a marked, linear slowdown with age across the remainder of the lifespan. From artists’ thirties to their eighties, tempo decreased by almost 10 bpm, averaging around 2 bpm per decade. This decrease aligns with the slowing-with-age hypothesis and mirrors rates of decline observed in studies of spontaneous motor tempo (SMT) and gait speed. Our findings highlight a significant gap in understanding of creative performance across the lifespan, particularly the role of age as a mediating factor in musical tempo. Moreover, that a discernible decrease in tempo is apparent even in commercial recordings further emphasizes the inescapable connection between dynamics of motor behavior and timing of musical performance.

## INTRODUCTION

Tempo is an essential characteristic of music that can be manipulated to express different emotions (Eerola & Vuoskoski, 2013), musical styles (Li & Chan, 2011), and to build and release tension (Goodchild et al., 2016). From a listener’s perspective, tempo impacts perception of emotion (Webster & Weir, 2005), time perception and estimation (Droit-Volet et al., 2013; Pereira et al., 2022), and characteristics of music-induced body movement (Burger et al., 2014). Body movement and music are, in fact, intimately intertwined, especially in terms of timing-related factors (Luck & Toiviainen, 2012). Body movement is not only induced by music but required to both understand it (Leman & Maes, 2014) and of course create it. Typical speed of body movement, frequently indexed via spontaneous motor tempo (SMT) and gait speed, is known to decrease across the adult lifespan (McAuley, et al., 2006; Bohannon & Andrews, 2011). This slowdown, likely a result of increased muscle activation and slowdown of nerve conduction velocities (Chase et al., 1992), is consistent with the slowing-with-age hypothesis (Baudouin et al., 2004), and is thought to reflect a degradation of motor, neural, and perceptual processes (Hunter et al., 2001; Morgan et al., 1994; Salthouse, 1996), as well as changes in muscle mass, visuo-proprioceptive function, strength, and reaction time (Voelcker-Rehage, 2008).

Recent work provides tentative evidence for an intrinsic connection between this age-related slowdown and tempo of music created across the lifespan (Luck, 2024). Examination of almost 2000 songs released by 10 best-selling solo artists revealed that their mean album tempo has fallen across each of their careers by as much as one and a half standard deviations from their early 20s to their late 50s. This effect holds despite the presence of different songwriters, producers, executives, collaborators, etc. Furthermore, it reveals that commercial recordings can offer profound insights into a fundamental and understudied aspect of human functioning across the lifespan.

Nonetheless, while this work offers support for a connection between age and tempo of artistic output, it remains unclear how generalizable this effect is beyond a small sample of artists. In addition, while Luck (2024) identified a linear relationship between tempo of commercial recordings and artist age, a curvilinear relationship has been identified in studies of SMT and gait across the lifespan (McAuley et al., 2006; Bohannon & Andrews, 2011). To gain a more comprehensive picture of how performance tempo changes as a function of artist age, it is essential to study a broader range of musical output. Here, we address these issues by investigating 1) a larger sample of songs from 2) a larger number of artists with 3) greater style and geographic diversity. Specifically, we examine tempo of over sixteen thousand tracks released by more than 200 artists while utilizing a more sophisticated statistical approach. In light of the curvilinear nature of SMT and gait speed trajectory across the lifespan, we were interested in exploring whether a larger, more diverse sample would reveal a similar trajectory in commercial music recordings. We hypothesized that tempo of an artist’s commercial output would follow an overall downward trajectory across their adult lifespan. However, we further hypothesized that this relationship would take a curvilinear form similar to that observed in studies of more basic motor timing, with a small increase in tempo during artists’ early adulthood followed by a more pronounced decrease in tempo across the remainder of their lifespan. Identifying the age at which this decline became apparent was a key issue of interest.

## METHOD

We drew our corpus from the *Spotify 1.2M+ Songs* dataset (Figueroa, 2020). This dataset contains computationally-extracted values for a wide range of musical features, including tempo. All features are based on Spotify’s own algorithms, and as such their precise definition and calculation are unknown. However, Spotify’s tempo feature is widely used in the research community (Duman et al., 2022; Gulmatico et al., 2022; Al-Beitawi et al., 2020) and serves as a standard proxy. The dataset contains 850,944 unique songs from 165,365 artists. We compiled our dataset by downloading the entire catalog from MusicBrainz and then querying Spotify’s API for each album’s Universal Product Code (UPC). This process enabled the retrieval of album information, which was further complemented by obtaining track details for each album through subsequent queries to the Spotify API (Figueroa, 2020). To ensure accurate data analysis, data were thoroughly cleaned, and a trimming process was carried out prior to the data modelling phase.

Since our aim was to examine effect of artist age on track tempo, we developed a range of exclusion criteria concerning artist age and career length, album type, and track type. We excluded artists with careers spanning less than 20 years; artists with fewer than 3 albums; and artists whose birth year could not be identified. We further excluded all music released before 1955; recordings identified as being in the classical genre; recordings identified as being recorded in live performance contexts (i.e., concerts); unreleased tracks and rarities; posthumous albums; compilations, greatest hits, collections, and ‘best of’ albums; tracks with featured artists; and tracks with speechiness values > 0.33 (i.e., tracks likely to be podcasts, interviews, etc.)^1^. To further limit extramusical influences on song characteristics, soundtrack albums, Christmas albums, remix albums, and mixtapes were also excluded. All songs appearing on multiple albums were retained only in the album on which they first appeared. In addition, we excluded any track with missing information about the release year or tempo, and tracks either less than 1 min or greater than 10 min in length.

Our final corpus contained 14,556 tracks released by 207 artists between 1956 and 2020. Average song length was 3.95 min (*SD* = 1.36). To operationalize artist age, we subtracted artist birth year from track release year. In cases where the artist was a band, we used the lead singer or lead instrumentalist’s birth year. Average artist age was 65.09 (*SD* = 13.07). The final dataset included artists from a diverse range of genres spanning Rock (e.g., Pearl Jam, Green Day, Alice Cooper), Pop (e.g., Kylie Minogue, Usher, Cyndi Lauper, Barbra Streisand), Country & Folk (e.g., Dolly Parton, Willie Nelson), Jazz & Blues (e.g., Luther Allison, Kenny G), Electronic and Dance (e.g., Paul Kalkbrenner), Rap & Hip Hop (e.g., Tech N9ne), Metal (e.g., Judas Priest, Disturbed), Latin (e.g., El Gran Combo De Puerto Rico, Juan Gabriel), and more.

### Analysis

Analyses were carried out in R using the *lme4* package (Bates et al., 2015) for Linear Mixed Models (LMMs), and *gamm4* (Wood & Scheipl, 2020) for Generalized Additive Mixed Models (GAMMs). GAMMs can be considered an extension of the Generalized Mixed Models that incorporate smooth, non-linear functions of predictors that allow for more flexible modelling of complex relationships between variables (Wood, 2017). The 95% Confidence Intervals for the estimates were computed using a bootstrap method (*N* = 10000 simulations). The Minimum Detectable Effect Sizes (MDEF) of the LMMs were computed using a Monte Carlo simulation approach (*N* = 2000 simulations) via the *simr* package (Green & MacLeod, 2016).

## RESULTS

### GAMM Model 1

We initially predicted track tempo (in bpm) as a non-linear function (i.e., smooth term) of artist age. Moreover, as we were aware of a positive correlation between release year and tempo (Léveillé Gauvin, 2018), we also added a non-linear effect of release year on tempo to control for this effect. Artists were modelled as random intercepts, thus allowing each artist to retain their baseline tempo level. This approach was crucial because it allowed us to appreciate and account for individual differences between artists. By taking these differences into account, we could make more generalizable comparisons and understand how age might influence changes in tempo beyond the artist’s natural tendency. Restricted Maximum Likelihood (REML) was used to estimate the smoothing parameter (i.e., λ)^2^ (Wood et al., 2016). This parameter determines the smoothness and flexibility of the fitted curve, balancing the model’s fit to the data against its complexity. In sum, the GAMM was built with the formula:

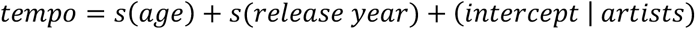

As expected, both terms reached statistical significance (*p*_year_ = .009; *p*_age_ = .013^3^).

Inspection of the predicted tempo (shown in Fig. 1) and effective degrees of freedom^4^ (*EDF*_age_ = 3.81) suggest that the relationship between age and tempo might deviate from being linear. The relationship between release year and tempo, however, is indistinguishable from a linear trend (*EDF*_year_ = 1.00)^5^. This suggests that the effect of age on tempo varies in a complex manner across artists’ lifespans, beyond what could be captured by a simple linear model.

**Figure 1.**
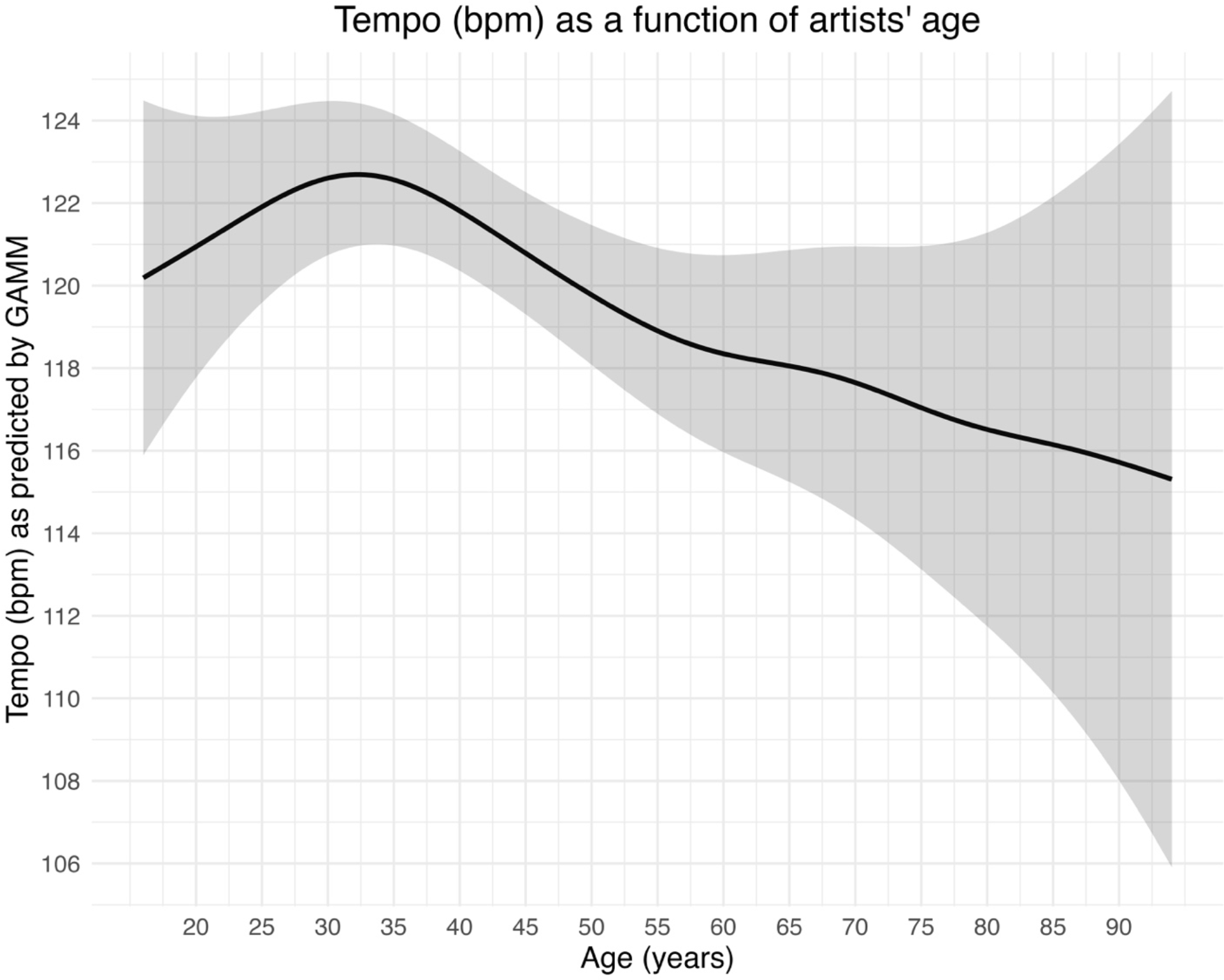
Tempo (bpm) as a function of artists’ age *Note*. The upper and lower confidence intervals are added at 2 standard errors above and below the mean.

### GAMM Model 2

Given variability in artists’ age-related tempo profile identified by Luck (2024), in a second step we examined whether the effect of age might differ significantly between artists. Consequently, a second more complex model was constructed in which artists were modelled as both random intercepts and slopes. Unlike Model 1, Model 2 permitted the effect of age on tempo to vary between artists. Specifically, in Model 2 we computed a unique curve for each artist, reflecting how each artist’s tempo changes differently over time. To build this model, we utilized the formula:

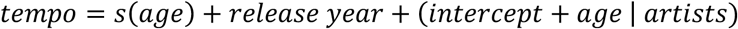

To test whether the complexity added in Model 2 improved or worsened data fit, a model comparison was performed (Rodgers, 2010) by inspecting the Akaike Information Criterion (AIC) (Pedersen et al., 2019), Bayesian Information Criteria (BIC), and Bayes Factor (Ward, 2008)^6^. Each of these metrics indicated a preference for Model 1, which included only random intercepts, demonstrating a better fit to the data [Table 1].

**Table 1.**
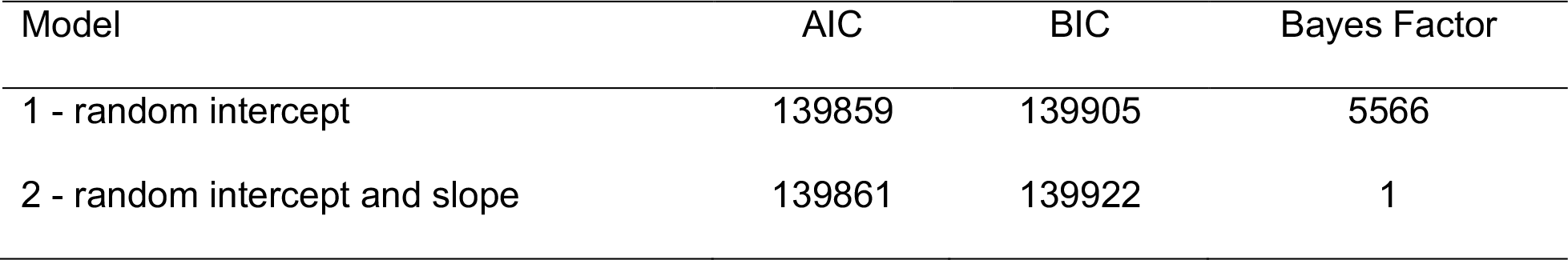
Model comparison.

| Model | AIC | BIC | Bayes Factor |
| --- | --- | --- | --- |
| 1 - random intercept | 139859 | 139905 | 5566 |
| 2 - random intercept and slope | 139861 | 139922 | 1 |

In summary, the increased values of AIC and BIC for Model 2, coupled with the Bayes Factor clearly favoring Model 1, indicate that the additional complexity introduced by the random slopes in Model 2 actually worsens the fit rather than improving it. For this reason, we retained the Model 1 configuration for subsequent analyses. Notably, this result implies that the effect of age on tempo is relatively consistent across artists. That is, the variability between the curves (i.e., how each artist changes their tempo over time) is relatively small, and the trajectories across artists are reasonably similar to each other when considering the non-linear effect of age.

Figure 1 shows tempo as a function of artist age. It can be seen that tempo increases slightly from artists’ mid-teens to early thirties, after which it decreases linearly over the rest of their lives. By the age of 50, tempo has dropped below mid-teen levels. By the age of 70, tempo has decreased by twice as much. By the age of 90, tempo has decreased three-fold.

### Analysis of the linear trends (LMMs)

To better assess the slopes of the increasing and decreasing trends, we took two further steps. First, we ascertained the linearity of the trends (i.e., the *EDF* values) by rerunning two instances of Model 1: One for the values of age < 30 (*EDF* = 1.00) and one for the values > 30 (*EDF* = 1.84). Finally, we ran a Linear Mixed Model (LMM) for each trend. The formula was identical to that of Model 1. The LMM for the increasing trend (i.e., age < 30; *N*_tracks_ = 2 237; *N*_artists_ = 96) signalled a positive relationship between age and tempo, *b* = 0.50, 95% CI [0.01,1.01], *SE* = 0.25, *p* = .043. The release year effect was not significant, *b* = −0.01, 95% CI [−0.20,0.18], *SE* = 0.09, *p* = .911. The LMM for the decreasing trend (i.e., age > 30; *N*_tracks_ = 12 023; *N*_artists_ = 204) indicated a significant negative relationship, *b* = −0.18, 95% CI [−0.29, −0.07], *SE* = 0.05, *p* = .001. In this model, the effect of the release year was significant, *b* = 0.16, 95% CI [0.05,0.27], *SE* = 0.06, *p* = .006.

This finding suggests that, after age 30, musicians’ tempo slows inexorably by approximately 1.8 beats per minute for every decade that passes.

A sensitivity analysis was conducted using a Monte Carlo simulation approach (Green & MacLeod, 2016) to evaluate the Minimum Detectable Effect Size (MDES) at power of 50% for the two models. In the case of the increasing trend model, the analysis determined that an effect size of *b* = 0.49 could be detected at power = 50.80%, 95% CI [48.58,53.01]. The effect we found could be detected at power = 52.45%, 95% CI [50.23,54.66]. As for the decreasing trend model, an effect size of *b* = −0.12 could be detected at power 51.10%, 95% CI [48.88,53.31], whereas our effect has power = 88.75%, 95% CI [87.28, 90.10].

## DISCUSSION

We investigated the effect of artist age on music performance tempo by examining computationally-extracted tempo values from a diverse range of commercial recordings. We utilized GAMMs and LMMs to partial out effects of artist age from more general effects of time (i.e., changing trends and other market forces) on performance tempo. Our analyses revealed a slight tempo increase from artists’ mid-teens to age 30, followed by a continuous, linear slowdown across the remainder of their lifespan. From the age of 30 to 80, tempo of musical output decreased linearly by about 10 bpm, averaging around 2 bpm per decade. This effect was independent of broader musical trends and resembles rates of decline observed in studies of spontaneous motor tempo (SMT) and gait speed.

Our results thus support the idea that changes in general motor competence across the lifespan, as evidenced in studies of SMT and gait speed, have a tangible impact on tempo of commercial music recordings. Moreover, the curvilinear nature of the relationship between music performance tempo and age identified here is closer to that of general indices of motor competence compared to the linear decline in tempo observed by Luck (2024). This is likely due to the larger, more diverse sample analyzed here affording greater detail and an order of magnitude more raw data points.

Despite the clear effects observed, there are several limitations that might be addressed in future work. First, the use of Spotify’s own tempo extraction algorithm resulted in a lack of clarity as to precisely how tempo was computed. Utilizing a non-black box approach in the future, if and where feasible, would afford greater transparency and understanding of results.

Second, in cases where the artist was a band or ensemble, operationalizing age as that of the lead singer or instrumentalist might not have provided the most accurate estimate of group age. In principle, a better, though considerably more time-consuming, approach might be to compute average age of all group members. However, this strategy would come at the cost of excluding a significant portion of the sampled artists. For instance, the lineup of many bands changes over time. Moreover, a large number of artists rely on session musicians for their recordings. Consequently, it will likely be challenging to identify the exact band composition of each album, even more so for each song. Furthermore, the birth year of many session musicians, who often work behind the scenes and contribute to recordings without being officially associated with a band or social artist, is not always easily available.

Third, our efforts to exclude tracks in order to minimize extramusical influences were not without their difficulties. For instance, some exclusions were based solely on names and nomenclature; a Christmas album title that did not contain the words “Christmas”, “Xmas” or similar would have been missed. In the same vein, we excluded classical music via typical related words (sonata, concerto, etc., and many names of classical composers). However, since track genre is not specified in the dataset, some lesser-known classical musicians/composers might have slipped through. Ideally, in subsequent studies, one might build one’s own dataset utilizing stricter criteria.

Fourth, we did not examine effects of musical genre on tempo slowdown across the lifespan. Future studies should also include this as a factor, if possible, in case, for example, rock musicians slow down faster or with a different trajectory than, say, pop artists, and so on.

Finally, we deliberately excluded live recordings from our sample since live contexts introduce additional confounding factors – click tracks to help set and maintain tempo and/or effects of heightened physiological arousal, for instance – likely to influence tempo. It could be valuable to investigate live recordings to see if similar effects of age on performance tempo can be observed.

From a creative perspective, our results have implications concerning the degree to which an artist’s age might inadvertently shape characteristics of their musical output. As artists age, they will become increasingly limited in terms of general motor competence, restricting the tempi at which they are likely to choose, or are indeed able, to perform. Considering how tempo is known to affect listeners’ experience, these effects have the potential to impact factors such as listeners’ perception of expression (Webster & Weir, 2005), their level of physiological arousal (Lundqvist et al., 2009), and characteristics of their music-induced body movement (Burger et al., 2014).

## Conclusions

We found that tempo of an artist’s musical output tends to decrease by about 2 bpm per decade from age 30 onwards. This decline likely results from degradation in general motor capacity since it mirrors concomitant degradation in SMT and gait speed. Moreover, this slowdown across the lifespan will likely impact how audiences experience or respond to artists’ output at different points in their careers. In short, our results reveal a notable gap in how we understand creative performance across the lifespan. In particular, we highlight the role of age as a mediating factor in music performance tempo. That an age-driven decrease in motor competence is apparent even in tempi of commercial recordings, which themselves are likely subject to a myriad of other influences, underscores the unavoidable link between timing of motor behavior and that of musical performance.

## Declarations

### Ethical approval

This study analyzed publicly available data obtained from Spotify, which does not involve personal or sensitive information. Consequently, the need for ethics approval was waived, as the project exclusively utilized data that are publicly accessible and do not infringe on individual privacy.

### Data Availability

The dataset analyzed during the current study is publicly available here: https://www.kaggle.com/datasets/rodolfofigueroa/spotify-12m-songs.

The R code was submitted via Manuscript Central and, at present, it is only available for review purposes.

## Footnotes

1 According to Spotify for Developers, speechiness “detects the presence of spoken words in a track. The more exclusively speech-like the recording (e.g. talk show, audio book, poetry), the closer to 1.0 the attribute value. Values above 0.66 describe tracks that are probably made entirely of spoken words. Values between 0.33 and 0.66 describe tracks that may contain both music and speech, either in sections or layered, including such cases as rap music. Values below 0.33 most likely represent music and other non-speech-like tracks.” (Spotify for Developers, 2024).

2 In the context of GAMMs, lambda (λ) acts as a penalization factor, controlling the trade-off between model complexity and fit accuracy. Higher λ values lead to greater penalization for complexity, resulting in more linear representations. Conversely, lower λ values allow for more detailed (non-linear) fits but risk overfitting. This mechanism ensures the model remains both accurate and generalizable.

3 Note that in the GAMM context, the null hypothesis is that the function is a flat line, i.e., the equivalent of testing the null hypothesis that the coefficient is 0 in a linear model.

4 The effective degrees of freedom (EDF) derived from Generalised Additive Models serve as an indicative measure of the non-linearity in the relationships. An EDF value of 1 signifies a linear relationship. When the EDF is between 1 and 2, the relationship is mildly non-linear. Conversely, an EDF > 2 suggests a pronounced non-linear relationship (Zuur et al., 2009).

5 When refitting the model by conceiving the release year as a linear term, its estimate was *b* = 0.14, 95% CI [0.03,0.24], SE = .05, *p* = .009.

6 AIC and BIC are rooted in information theory. They are designed to provide a measure of the relative quality of statistical models for a given set of data. They balance model complexity against the goodness of fit, essentially penalizing models that overfit by adding unnecessary parameters. Therefore, when comparing models, those with lower AIC or BIC values are preferred, indicating a more efficient balance between simplicity and explanatory power. On the other hand, the Bayes Factor provides a direct comparison between two models, quantifying the evidence in favor of one model over another. As such, higher BF values indicate stronger evidence for the preferred model (Ward, 2008; Zhang et al., 2023).

## References

Al-Beitawi, Z., Salehan, M., & Zhang, S. (2020). What Makes a Song Trend? Cluster Analysis of Musical Attributes for Spotify Top Trending Songs. Journal of Marketing Development and Competitiveness, 14(3). Retrieved from https://articlearchives.co/index.php/JMDC/article/view/4610

Bates, D., Mächler, M., Bolker, B., & Walker, S. (2015). Fitting Linear Mixed-Effects Models Using lme4. Journal of Statistical Software, 67(1). 10.18637/jss.v067.i01

Baudouin, A., Vanneste, S., & Isingrini, M. (2004). Age-Related Cognitive Slowing: The Role of Spontaneous Tempo and Processing Speed. Experimental Aging Research, 30(3), 225– 239. 10.1080/03610730490447831

Bohannon, R., & Andrews, A. (2011). Normal walking speed: a descriptive meta-analysis. Physiotherapy, 97(3), 182–9. 10.1016/j.physio.2010.12.004.

Burger, B., Thompson, M. R., Luck, G., Saarikallio, S., & Toiviainen, P. (2014). Hunting for the beat in the body: On period and phase locking in music-induced movement. Frontiers in Human Neuroscience, 8(903).

Chase, M. H., Engelhardt, J. K., Adinolfi, A. M., & Chirwa, S. S. (1992) Age-dependent changes in cat masseter nerve: An electrophysiological and morphological study. Brain Research, 586(2), 279–288.

Droit-Volet, S., Ramos, D., Bueno, J. L. O., & Bigand, E. (2013). Music, emotion, and time perception: The influence of subjective emotional valence and arousal? Frontiers in Psychology, 4. 10.3389/fpsyg.2013.00417

Duman, D., Neto, P., Mavrolampados, A., Toiviainen, P., & Luck, G. (2022). Music we move to: Spotify audio features and reasons for listening. PLOS ONE, 17(9), e0275228. 10.1371/journal.pone.0275228

Eerola, T., & Vuoskoski, J. K. (2013). A Review of Music and Emotion Studies: Approaches, Emotion Models, and Stimuli. Music Perception, 30(3), 307–340. 10.1525/mp.2012.30.3.307

Figueroa, R. (2020). Spotify 1.2M+ Songs. Available at: https://www.kaggle.com/datasets/rodolfofigueroa/spotify-12m-songs

Goodchild, M., Gingras, B., & McAdams, S. (2016). Analysis, Performance, and Tension Perception of an Unmeasured Prelude for Harpsichord. Music Perception, 34(1), 1–20. 10.1525/mp.2016.34.1.1.

Green, P., & MacLeod, C. J. (2016). SIMR: An R package for power analysis of generalized linear mixed models by simulation. Methods in Ecology and Evolution, 7(4), 493–498. 10.1111/2041-210X.12504

Gulmatico, J. S., Susa, J. A. B., Malbog, M. A. F., Acoba, A., Nipas, M. D., & Mindoro, J. N. (2022). SpotiPred: A Machine Learning Approach Prediction of Spotify Music Popularity by Audio Features., in 2022 Second International Conference on Power, Control and Computing Technologies (ICPC2T), (Raipur, India: IEEE), 1–5. doi: 10.1109/ICPC2T53885.2022.9776765

Hunter, S. K., Thompson, M. W., & Adams, R. D. (2001). Reaction Time, Strength, and Physical Activity in Women Aged 20–89 Years. Journal of Aging and Physical Activity, 9(1), 32– 42. 10.1123/japa.9.1.32

Leman, M., & Maes, P.-J. (2015). The Role of Embodiment in the Perception of Music. Empirical Musicology Review, 9(3–4), 236–246. 10.18061/emr.v9i3-4.4498

Léveillé Gauvin, H. (2018). Drawing listener attention in popular music: Testing five musical features arising from the theory of attention economy. Musicae Scientiae, 22(3), 291– 304. 10.1177/1029864917698010

Li, T. LH., & Chan, A. B. (2011). Genre classification and the invariance of MFCC features to Key and Tempo. In: International Conference on MultiMedia Modeling (pp. 317–217). Berlin, Heidelberg; Springer.

Lundqvist, L.-O., Carlsson, F., Hilmersson, P., & Juslin, P. N. (2009). Emotional responses to music: Experience, expression, and physiology. Psychology of Music, 37(1), 61–90. 10.1177/0305735607086048

Luck, G. (2024). Neurobiological slowdown in later life manifests in tempo of popular music. bioRxiv preprint doi: 10.1101/2024.02.06.579086

Luck, G., & Toiviainen, P. (2012). Movement and musical expression. In A. R. Brown (Ed.) Sound Musicianship: Understanding the Crafts of Music. Cambridge Scholars Publishing; Newcastle upon Tyne, UK. 167–177.

McAuley, J. D., Jones, M. R., Holub, S., Johnston, H. M., & Miller, N. S. (2006). The time of our lives: Life span development of timing and event tracking. Journal of Experimental Psychology: General, 135(3), 348–367. 10.1037/0096-3445.135.3.348

Morgan, M., Phillips, J. G., Bradshaw, J. L., Mattingley, J. B., Iansek, R., & Bradshaw, J. A. (1994). Age-related motor slowness: Simply strategic? Journal of Gerontology, 49(3), M133–M139. 10.1093/geronj/49.3.M133

Pedersen, E. J., Miller, D. L., Simpson, G. L., & Ross, N. (2019). Hierarchical generalized additive models in ecology: An introduction with mgcv. PeerJ, 7, e6876. 10.7717/peerj.6876

Pereira, L. A. S., Ramos, D., & Bueno, J. L. O. (2022). The influence of different musical modes and tempi on time perception. Acta Psychologica, 229, 103701. 10.1016/j.actpsy.2022.103701

Rodgers, J. L. (2010). The epistemology of mathematical and statistical modeling: A quiet methodological revolution. American Psychologist, 65(1), 1–12. 10.1037/a0018326

Salthouse, T. A. (1996). The processing-speed theory of adult age differences in cognition. Psychological Review, 103(3), 403–428. 10.1037/0033-295X.103.3.403.

Spotify for Developers (2024). Get Track’s Audio Features. Available at: https://developer.spotify.com/documentation/web-api/reference/get-audio-features (Accessed March 16, 2024).

Voelcker-Rehage, C. (2008). Motor-skill learning in older adults — A review of studies on agerelated differences. European Review of Aging and Physical Activity, 5(1), 5–16. 10.1007/s11556-008-0030-9

Ward, E. J. (2008). A review and comparison of four commonly used Bayesian and maximum likelihood model selection tools. Ecological Modelling 211, 1–10. 10.1016/j.ecolmodel.2007.10.030

Webster, G. D., & Weir, C. G. (2005). Emotional Responses to Music: Interactive Effects of Mode, Texture, and Tempo. Motivation and Emotion, 29(1), 19– 39. 10.1007/s11031-005-4414-0

Wood, S. (2017). Generalized Additive Models: An Introduction with R., 2nd Ed. Chapman and Hall/CRC. 10.1201/9781315370279

Wood, S. N., Pya, N., & Säfken, B. (2016). Smoothing Parameter and Model Selection for General Smooth Models. Journal of the American Statistical Association 111, 1548– 1563. 10.1080/01621459.2016.1180986

Wood, S., & Scheipl, F. (2020). gamm4: Generalized Additive Mixed Models using “mgcv” and “lme4.” Available at: https://cran.r-project.org/web/packages/gamm4/index.html

Zhang, J., Yang, Y., & Ding, J. (2023). Information criteria for model selection. WIREs Computational Statistics, 15(5), e1607. 10.1002/wics.1607

Zuur, A. F., Ieno, E. N., Walker, N., Saveliev, A. A., & Smith, G. M. (2009). Mixed effects models and extensions in ecology with R. New York, NY: Springer New York. 10.1007/978-0-387-87458-6

